# JAK1/2 Inhibition Delays Cachexia and Improves Survival through Increased Food Intake

**DOI:** 10.1101/2025.10.21.683287

**Authors:** Ezequiel Dantas, Anirudh Murthy, Jeshua Kim, Tanvir Ahmed, Mujmmail Ahmed, Tiffany Perrier, Shakti Ramsamooj, Isaac Nathoo, Lucas Kniess Debarba, Andre Lima Queiroz, Shiri Li, Baran Ersoy, Miriam Ferrer, Ido Goldstein, Jonathan Gao, Tiffany Lam, Matthew Nagler, Murtaza Malbari, Nasser Altorki, Eduardo Cararo Lopes, Maria Gomez Jenkins, Trishna Das, Mariam Jamal-Hanjani, Eileen White, Tobias Janowitz, Marcus D Goncalves

## Abstract

Lung cancer is the leading cause of cancer-related death and is frequently accompanied by reduced food intake and cachexia, a debilitating syndrome characterized by weight loss and skeletal muscle wasting. We sought to identify contributors to cachexia using a murine model of lung cancer that reproduces key features of this syndrome. A multiplex cytokine screening approach, integrated with western blot and transcriptomic analyses, identified tumor-derived inflammatory mediators and downstream signaling pathways associated with cachexia. Notably, IL-6 superfamily members were elevated in the tumor and plasma of mice and patients with cachexia. The JAK-STAT3 signaling was upregulated in liver and skeletal muscle, driving the acute phase response and impairing lipid metabolism. Pharmacologic inhibition of JAK1/2 with ruxolitinib improved body weight, fat mass, and overall survival without altering tumor burden. These effects were driven primarily by blunted hypothalamic leptin receptor signaling, which increased food intake early in the disease course. In the liver, JAK inhibition reduced STAT3 activity, restored fatty acid oxidation, and decreased the production of acute-phase proteins. These findings support JAK inhibition as a therapeutic strategy for lung cancer-associated cachexia.

**Statement of Significance:** Cancer cachexia is a lethal complication of lung cancer that lacks effective treatment. We show that JAK inhibition by ruxolitinib restores weight, fat mass, and prolongs survival in murine models of lung cancer. These effects were independent of tumor burden, underscoring the relevance of addressing cachexia to improve survival in cancer patients and supporting clinical testing of JAK inhibition for cancer cachexia

## Introduction

Lung cancer is the leading cause of cancer death among both men and women^1^. Most patients with lung cancer die with reduced food intake (anorexia) and cachexia, a systemic wasting syndrome characterized by the loss of body weight and skeletal muscle mass^2^. Anorexia and cachexia frequently co-occur and both contribute to involuntary weight loss, making it difficult to distinguish their individual contributions using current clinical diagnostic criteria. The current diagnostic criteria for cachexia includes loss of 5% or more of their body weight over 6 months, or loss of ≥2% of body weight over 6 months when the body mass index (BMI) is already low (<20 kg/m^2^)^3^. This degree of weight loss is common in patients with lung cancer and correlates with reduced quality of life, increased treatment complications, and worsened mortality^4–6,7–9^. Therefore, treating or preventing cachexia is predicted to improve quality of life, prolong the duration of anti-cancer therapy, and elongate overall survival.

Tumor molecular features contribute to clinical outcomes and the development of cachexia. Loss of STK11/LKB1 activity is prevalent in *KRAS*-mutant lung cancer and promotes aggressive and treatment-resistant tumors^10,11^. Somatic loss of function mutations in *LKB1* correlate with the presence of cachexia in newly diagnosed patients with lung cancer^9,12^, and these phenotypes are recapitulated in mice with tumors induced by *Kras* activating mutations and loss of function *Lkb1* alleles in the lung (KL mice)^13,14^. KL mice develop aggressive, adenocarcinoma and adenosquamous tumors that are resistant to chemotherapy and immunotherapy^13,15^. These mice also develop a cancer-induced anorexia and cachexia syndrome (CACS) with features consistent with the human condition such as weight loss, skeletal muscle and adipose atrophy, anorexia, and reduced physical performance^14,16^.

Mouse models have greatly aided in the identification of cachexia mediators. Several groups have found that tumor-secreted factors play pathophysiologic roles in weight loss. For example, inhibition of IL-6 and PTHrP can prevent cachexia in mice bearing C26- and LLC-allografts, respectively, which secrete these factors^17,18^. Treating cachexia in autochthonous genetic mouse models has proven to be more challenging; however, we recently showed that a combination of therapies targeting appetite stimulation and Activin A signaling can reverse CACS in female KL mice^16^. Interestingly, this approach was not effective in male mice of the same background despite a similar degree of anorexia and equivalent levels of Activin A. The specific drivers of CACS in these male mice remain unknown.

In both humans and mice with cancer, cachexia correlates with elevated markers of chronic inflammation, including C-reactive protein (CRP) and cytokines such as tumor necrosis factor-alpha (TNF-α), interleukin-1 (IL-1), and IL-6^19^. Other circulating immune factors like GDF-15, Activin A, CCL2, and Lipocalin 2 are also upregulated and likely contribute to the systemic metabolic dysfunction that promotes cachexia. However, clinical studies using therapeutic antibodies against pro-inflammatory cytokines, like TNF-α or IL-6, have not shown significant benefit^20–23^, and studies targeting GDF-15 are ongoing^24,25^. Therefore, there is an urgent need to further understand the contribution of inflammation to cachexia during cancer development and progression.

Pro-inflammatory cytokines promote CACS in several ways. They can act centrally in the brain to disrupt normal appetite regulation and contribute to negative energy balance^19^. In the periphery, cytokines can bind directly to skeletal muscle and adipose tissue to initiate catabolic processes like proteolysis and lipolysis, respectively. Furthermore, some cytokines are known to induce the acute phase response (APR) in the liver, which enhances immunity, aids in pathogen clearance, and promotes tissue repair^26,27^. Activation of the APR is a poor prognostic factor in human cancer ^28–31^, especially in patients with lung cancer ^32^. The APR requires a significant change in hepatic metabolism to accommodate the large increase in de novo protein production and release by the liver ^33,34^. For example, many genes involved in redox processes and fatty acid oxidation (FAO) are downregulated in response to inflammatory stimuli like IL-6^35,36^.

In this study, we explore the relationships between cancer, pro-inflammatory cytokines, anorexia, and cachexia. We find that CACS is associated with increased production of IL-6 family members by the tumor, which subsequently accumulate in the serum, leading to activation of the pro-inflammatory JAK/STAT3 pathway in peripheral tissues. In the liver, JAK signaling triggers the APR and inhibits FAO via PPAR-α interactions, a pattern consistent with cachexia across various animal models and human samples. Systemic inhibition of JAK not only suppresses the APR and reactivates FAO pathways in the liver but also enhances food intake and fat mass by suppressing leptin sensing in the brain, thereby extending survival independently of tumor progression.

## Results

Following exposure to inhaled adenovirus carrying Cre recombinase, male KL mice develop the CACS phenotype characterized by the loss of body weight (Fig. 1A and B), fat mass (Fig. 1C), and lean mass (Fig. 1D). Since body weight is determined by the balance between energy intake and energy expenditure, we measured these components over time in mice with CACS. At 4 weeks after tumor induction, we observed declines in food intake and physical activity (Fig 1E, F,). These changes preceded reductions in body weight and changes in body composition (Fig. 1B, C, D). Further declines in food intake and activity were observed at 9 weeks after induction, when total energy expenditure was also significantly reduced (Fig 1G, H). These results suggest that in KL mice, weight loss and body composition changes are primarily driven by reduced food intake, rather than by an increase in energy demands.

**Figure 1.**
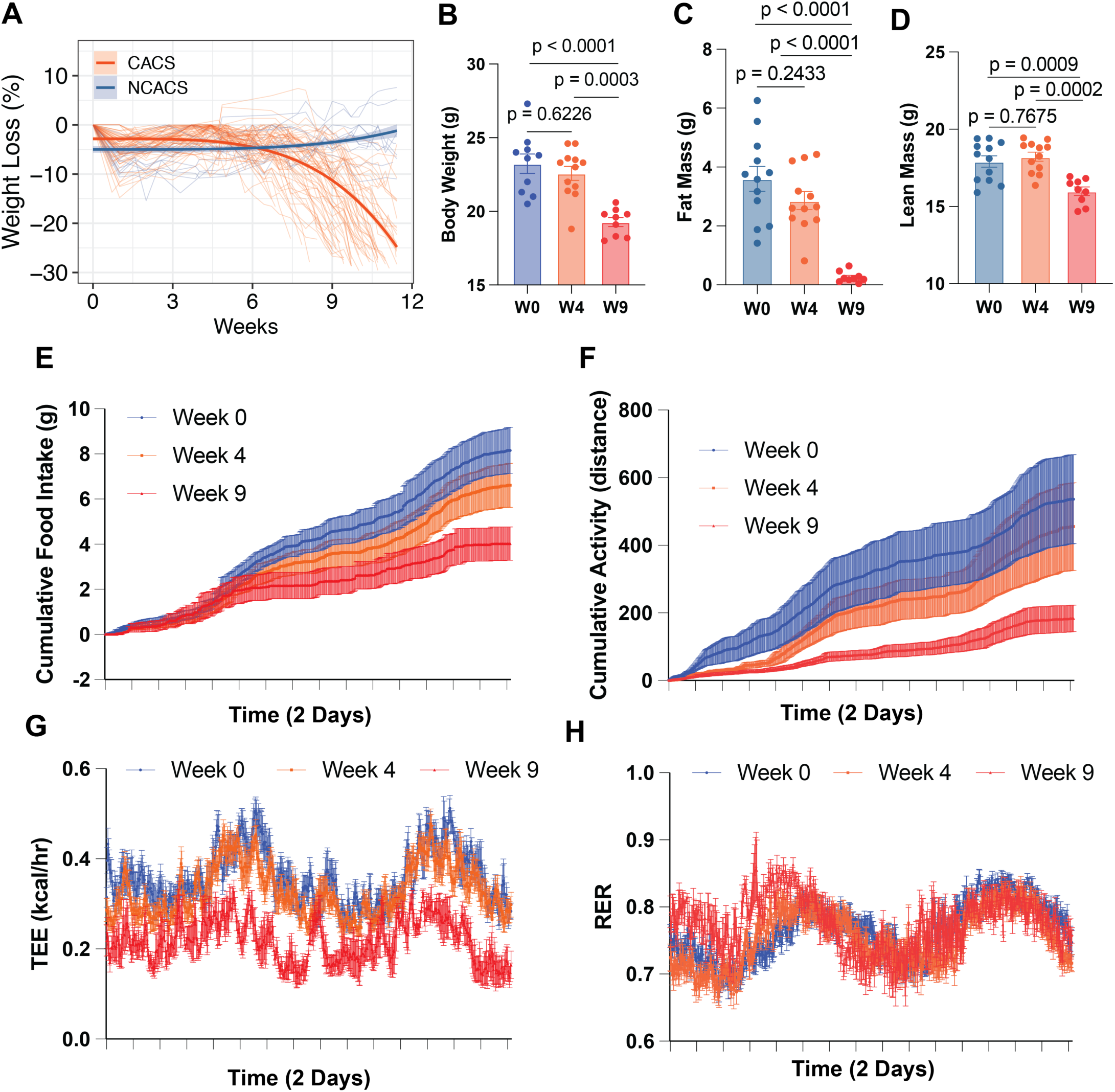
Cachexia is associated with anorexia and low total energy expenditure. **A.** Progression of weights normalized to peak from male KL mice (A). **B-D.** Body weight, fat mass and lean mass from KL mice before induction (W0) and after four (W4) and nine weeks (W9) after induction. E-H. Cumulative food intake in grams (E), Total Energy Expenditure (F), cumulative activity (G) and respiratory exchange ratio for the mice displayed in B, C and D. Comparisons in B, C and D were performed with one-way ANOVA followed by Tukey’s multiple comparisons test. Individual data points are independent biological replicates unless otherwise stated.

About 20% of KL mice maintain their body weight (referred to as NCACS), despite achieving similar amounts of tumor burden^14,16^. This incomplete penetrance offered us a unique opportunity to compare mice with and without CACS in a setting controlled for genetic and environmental exposures. To understand the determinants that drive weight loss, we isolated tumors from mice with and without CACS and performed transcriptome analysis using RNA-Seq to identify potential mediators. Using Ingenuity Pathway Analysis, we found that tumors isolated from mice with CACS had significant enrichment in pathways related to inflammation like “Neutrophil Extracellular Trap Signaling”, “Macrophage Alternative Activation Signaling”, “Toll-like Receptor Signaling” and IL-6, IL-17 and IL-33 Signaling (Fig 2A). To validate these results in patients with lung cancer, we mined the National Cancer Institute’s Clinical Proteomic Tumor Analysis Consortium (CPTAC) lung adenocarcinoma (LUAD) dataset^37^, for which there is RNA-Seq, proteomics, phospho-proteomics and body mass index (BMI) available. This data allowed us to stratify patients by “low BMI” (<20 kg*m^-2^), a surrogate for cachexia^3^, or “high BMI” (>20 kg*m^-2^) to compare the tumor proteomic signatures. We found that patients with low BMI have tumors enriched in pathways related to inflammation, IL6/STAT3, TNF-alpha/NFKB, and interferon-alpha response (Fig. 2B) in agreement with the KL mice. Similarly, a gene set enrichment analysis (GSEA) of primary lung cancer RNA-Seq data from the TRAcking non-small cell lung Cancer Evolution through therapy (Rx) (TRACERx) study (ClinicalTrials.gov identifier: NCT01888601) showed that patients who develop cachexia at relapse after surgical resection also have enrichment in “Inflammatory response pathway” (Supp. Fig1. A)^38^.

**Figure 2.**
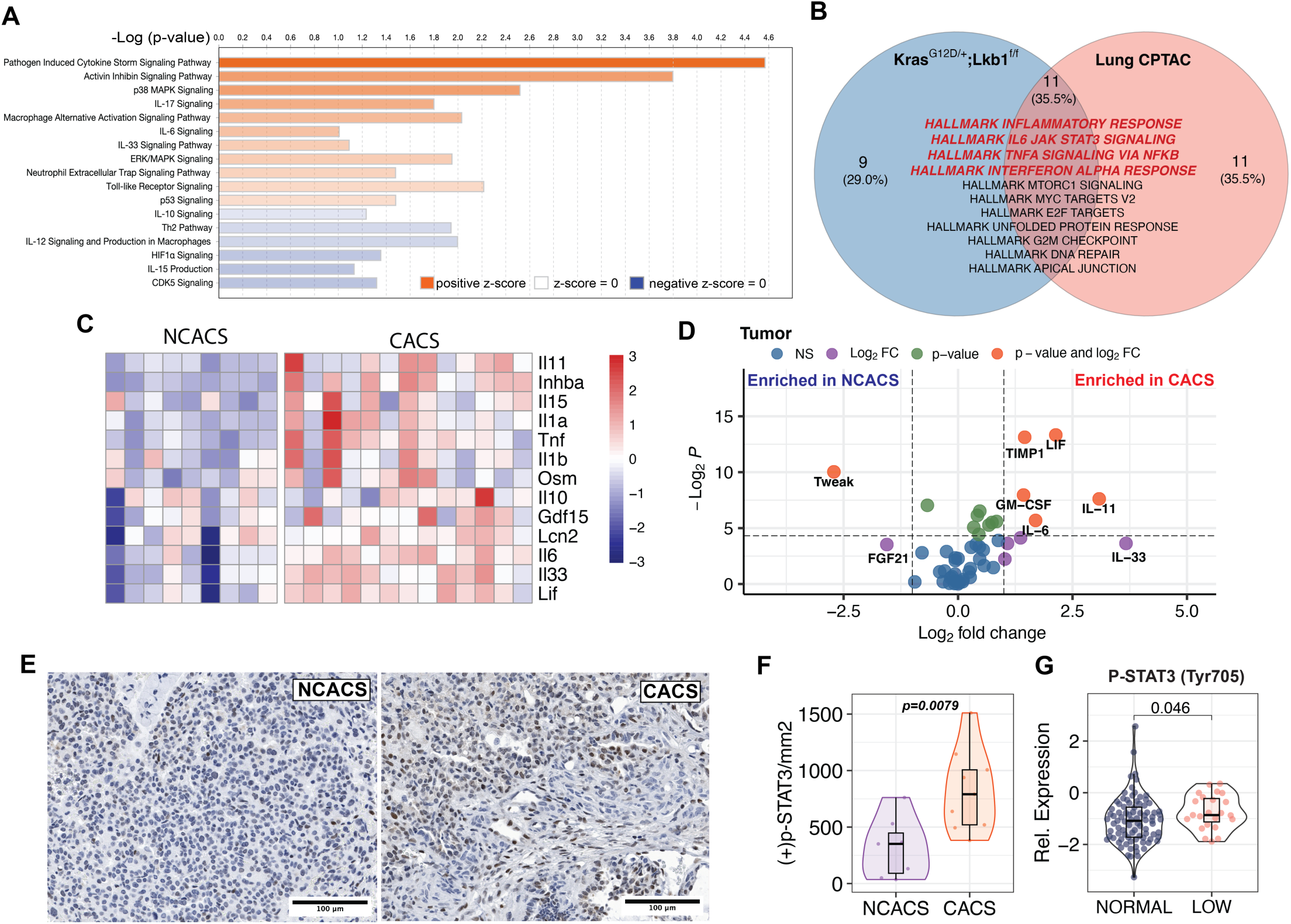
Cachexia is associated with a pro-inflammatory tumor microenvironment. **A.** Ingenuity Pathway Analysis (IPA) comparing tumor transcriptomics between CACS and NCACS mice. **B.** Venn-diagram showing the results from KL mice tumor transcriptomics Gene Set Enrichment Analysis (GSEA) comparing CACS to NCACS (blue) and tumor proteomics from patients with low and high body mass index (BMI) in the CPTAC cohort (red). **C.** Heatmap that shows the differential expression of cytokines relevant to cachexia in tumors from CACS and NCACS mice. **D.** Volcano plot comparing cytokine levels measured by Luminex in tumor lysates from CACS and NCACS. **E.** Immunohistochemistry (IHC) for p-STAT3 (Tyr705) in the KL tumors from CACS and NCACS mice. **F.** Quantification of the positive cells per square millimeter in the tumor p-STAT3 (Tyr705) IHC of CACS and NCACS KL mice. **G.** Relative expression of p-STAT3 (Tyr705) comparing tumor phospho-proteomics from patients with low and normal BMI in CPTAC cohort. Comparisons in D, F, G were done with Wilcoxon’s test. Comparisons in M and O were performed with one-way ANOVA followed by Tukey’s multiple comparisons test. Individual data points are independent biological replicates unless otherwise stated.

The genes driving the inflammatory GSEA in mice included many of those coding for cytokines and immune-related factors that have previously been linked to cachexia (Fig. 2C)^19^. We next performed a multi-analyte profiling of tumor lysates to assess the abundance of 45 cytokines and chemokines. This analysis identified significant enrichment in three IL-6 superfamily members (IL-6, LIF, IL-11) in the tumors of mice with CACS (Fig. 2D, and Supp. Fig. 1, B,C and D). In agreement with the elevated gene expression and increased abundance of IL-6 superfamily members, we found histologic evidence of increased JAK signaling in the tumors using immunohistochemistry against phosphorylated [residue] on STAT3 (Fig. E and F). Similarly, the phosphorylation of STAT3 at Tyr705 was found to be higher in the tumors from patients with low BMI from the CPTAC dataset (Fig. 2G). High expression of these IL-6 superfamily members in the tumors of patients with lung cancer correlates with poor overall survival, a feature consistent with cachexia (Supp. Fig. 1B, C, and D). In support, we also found increased levels of IL-6 in the plasma of patients with lung cancer from an independent cohort (Supp. Fig.1E). Of note, IL-11 was also elevated in the plasma of patients with lung cancer but the comparison between CACS and NCACS did not reach statistical significance (Supp. Fig.1F). These data support the correlation between inflammation, JAK signaling, and cachexia.

It is commonly believed that tumor-derived factors induce cachexia by exiting the tumor, circulating in the blood, and signaling to end-organs like skeletal muscle and liver. Therefore, we repeated the analyte profiling using serum from mice with and without CACS. Of the IL-6 family members found differentially expressed in the tumor, only IL-6 was significantly enriched in the serum, albeit at a relatively low absolute concentration (< 20 pg/mL) (Fig 2A). Additionally, IL-33 and IL-10, which can activate JAK/STAT3, were also found to be increased in the serum of CACS mice (Fig. 3A).

**Figure 3.**
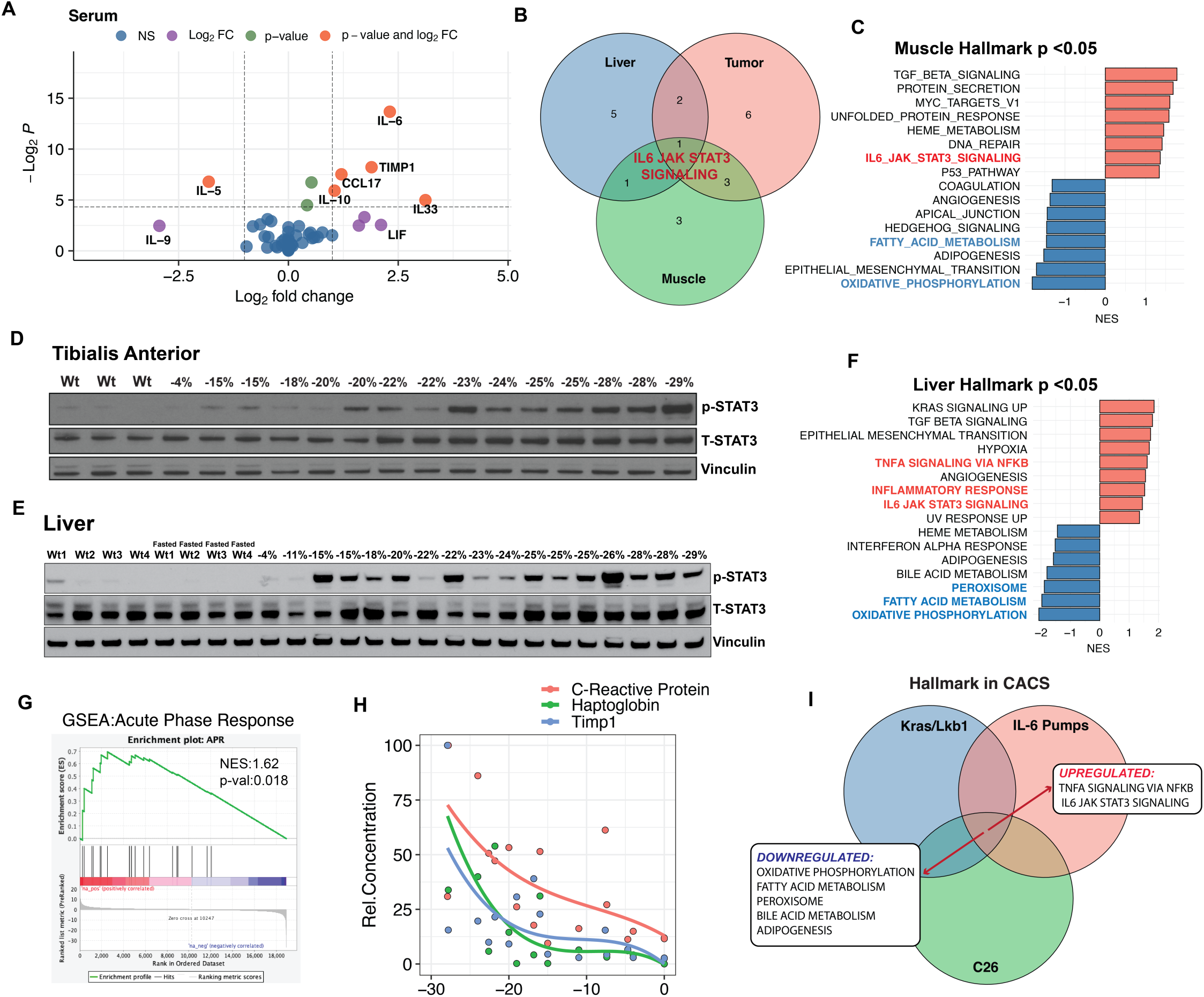
Cachexia is associated with systemic JAK activation. **A.** Volcano plot comparing cytokine levels measured by Luminex in plasma from CACS and NCACS KL mice. **B.** Venn-diagram with common GSEA results of the Hallmark database for liver, tumor and muscle. **C.** GSEA comparing muscle transcriptomics between CACS and NCACS KL mice using the Hallmark database. **D.** Western Blot (WB) for p-STAT3 (Tyr705), Total STAT3 and Vinculin in the tibialis anterior (TA) of non-tumor-bearing (WT) mice and KL mice with different stages of weight loss. **E.** GSEA comparing between livers of CACS and NCACS mice. **F.** Western Blot (WB) for p-STAT3 (Tyr705), Total STAT3 and Vinculin in liver of non-tumor-bearing (WT) mice, WT mice with 15 hours of food restriction, and KL mice with different stages of weight loss **G.** GSEA of acute phase reactants (APR) when comparing the liver transcriptomics of CACS and NCACS KL mice. **H.** Relative concentration of APR proteins (C-reactive protein, Haptoglobin, and TIMP1) in the serum of KL mice with different grades of weight loss. APR concentration was normalized to 1-100 to plot all the analytes in a single graph. **I.** Venn-diagram showing GSEA common results between liver transcriptomics from KL mice (CACS vs NCACS), C26 tumor-bearing mice (tumor vs PBS control), and mice implanted with IL-6 pumps (IL-6 pump vs PBS pump). Comparison in A was done with Wilcoxon’s test. Individual data points are independent biological replicates unless otherwise stated.

In agreement with elevated levels of IL-6 superfamily members, we found evidence for systemic JAK/STAT3 activation in several tissues from the KL mice with CACS. GSEAs of the tumor, liver, and skeletal muscle transcriptomics shared an enriched activation of the IL6/JAK/STAT3 pathway in mice with CACS (Fig. 3B). In the muscle, this pathway is one of the most differentially expressed in the mice with CACS, as compared to NCACS (Fig. 3C). The activation of the JAK pathway in the muscle was confirmed using western blot, which showed a clear positive correlation between STAT3 phosphorylation and weight loss (Fig. 3D).

The liver is not only a metabolic organ but also a key player in both innate and adaptive immunity. Hepatocytes and other liver cells express multiple cytokine receptors, including those for IL-6 family cytokines. This expression enables the liver to sense and respond to pathogens and inflammatory signals, integrating metabolic regulation with immune responses. In agreement with the results from skeletal muscle, we found a clear positive correlation between hepatic STAT3 phosphorylation (Tyr705) and weight loss in tumor-bearing mice, which did not occur in non-tumor-bearing mice or an overnight fast (Fig. 3E). Similarly, we observed a marked enrichment of inflammatory pathways in the cachexic transcriptome, including the TGF-beta, TNF-alpha, and STAT3 signaling pathways (Fig. 3F). These results suggest that the phenotype of the cachexic liver is markedly influenced by inflammatory factors during CACS.

The hepatic response to inflammation includes marked changes to protein and lipid metabolism. For example, pro-inflammatory stimuli are known to induce the APR in hepatocytes, which enhances the production of proteins that support immunity, aids in pathogen clearance, and promote tissue repair^26,27^. Indeed, the livers of mice with CACS showed a marked increase in the transcription of genes related to the APR response (Fig. 3G), and there was a rise in the serum concentration of APR proteins (*e.g., Haptoglobin*) during weight loss (Fig. 3H). Hepatic fat metabolism is also perturbed by inflammation, and we observed a suppression of several lipid metabolism pathways in the liver of mice with CACS (Fig. 3F), which agrees with previous reports of reduced peroxisome proliferator-activated receptor-α (PPAR-α) activity and ketogenesis in the cachexic liver^14,36^. These data led us to hypothesize that the rise in APR and suppression of PPAR-α related pathways are mechanistically linked to a common upstream mediator.

To investigate whether the metabolic changes observed in the cachexic liver are mediated by IL-6 superfamily members, we implanted mice with osmotic pumps containing IL-6 or PBS, as controls. The mice were allowed ad-libitum food intake for the first five days and then food was withheld for 24 hours to induce PPAR-α and ketogenesis (Supp. Fig 2A). In control mice, the fasting period reduced body weight (Supp. Fig 2B) and increased ketogenesis, as measured by a rise in serum levels of beta hydroxybutyrate (BHB, Supp. Fig.2C). Exposure to IL-6 significantly induced JAK activity in the liver (Supp. Fig.2D) and suppressed fasting ketogenesis, without altering the degree of weight loss (Supp. Fig.2B and C). Next, we performed RNA-Seq on the IL-6 and PBS-exposed liver tissue. The mice exposed to both IL-6 and fasting clustered independently from the fed and fasted mice exposed to PBS using unbiased principal component analysis (PCA) (Supp Fig. 2E), suggesting that IL-6 produces distinct transcriptomic changes compared to fasting alone. Using GSEA, we found that IL-6 enriched the liver in IL-6/JAK/STAT signaling and inflammation pathways, as compared to fasted PBS control mice (Supp. Fig.2F). IL-6 treatment was also sufficient to activate the expression of genes related to the APR (Supp. Fig. 2G) and suppress the expression of transcripts and pathways related to PPAR-α activity (Supp. Fig. 2H). To delineate the contribution of IL-6 family members to cachexia, we compared the GSEA results from the IL-6 pumps to those from the liver of KL mice and the liver of mice bearing allografts of the C26 cell lines, another commonly utilized cachexia model^39^. Activation of JAK and suppression of PPAR-α-related pathways were conserved amongst all data sets (Fig. 3I). Together, these data suggest that tumor-induced cytokines promote JAK/STAT3 activity in the liver, which facilitates a metabolic rewiring characterized by an increase in the APR and suppression of PPAR-α activity.

Therefore, we next sought to target JAK activity directly with pacritinib, an FDA-approved multi-kinase inhibitor that targets JAK2, a key kinase for the signaling of the IL-6 family receptors. Pacritinib was compounded into the chow at a concentration of 0.3%, per the manufacturer’s suggestion, and we performed pilot studies in non-tumor-bearing mice to confirm that this approach is safe (no change in food intake, activity, grooming habits, and weight) and yields similar drug exposures as a previously published dose of 100 mg/kg twice daily (data not shown)^40^. Therefore, we randomized mice to receive pacritinib or control at 4 weeks after induction, and overall survival was our primary outcome. Pacritinib tended to delay weight loss and improve the overall survival, without changes in tumor or skeletal muscle mass (Fig. 4 A, B and C, Supp. Fig.3 A and B). However, the effect was not statistically significant.

**Figure 4.**
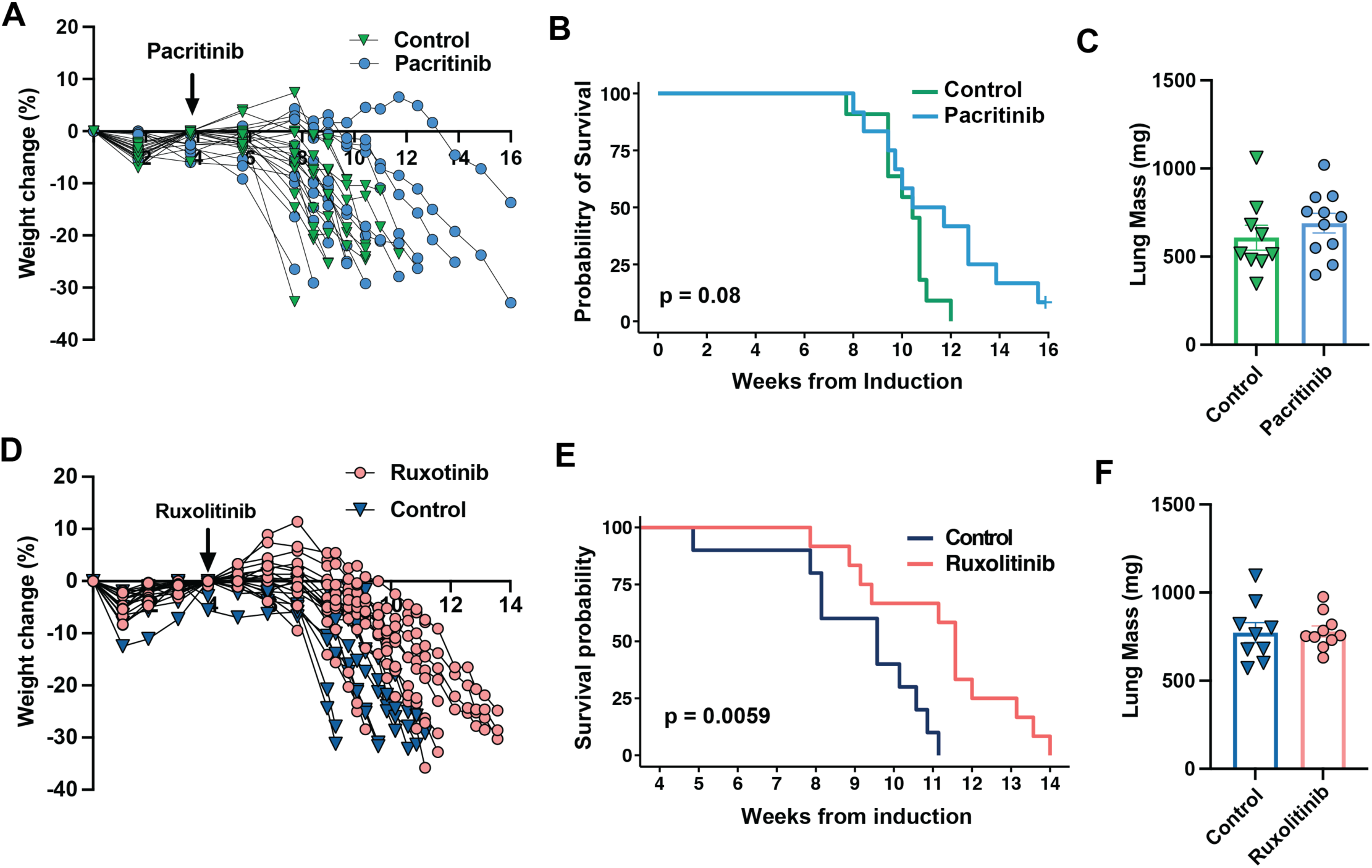
A. Jak1/2 inhibition prolongs survival in male mice with cachexia. **A.** Progression of weights normalized to peak from cachexic KL mice treated with Pacritinib. **B.** Kaplan-Meier plot with the probability of survival from cachexic KL mice treated with pacritinib. **C.** Lung mass of KL mice treated with pacritinib. **D.** Progression of weights normalized to peak from cachexic KL mice treated with ruxolitinib. **E.** Kaplan-Meier plot with the probability of survival from cachexic KL mice treated with ruxolitinib. **F.** Lung mass of KL mice treated with ruxolitinib at endpoint. Comparisons for the KM plots in B, and E were done with the Log-rank Mantel-Cox test. Comparisons in C and F were done using two-tailed Student’s t-test. Individual data points are independent biological replicates unless otherwise stated.

The partial response to pacritinib encouraged us to pursue an additional study using the FDA-approved drug, ruxolitinib, which inhibits both JAK1 and JAK2. Ruxolitinib was compounded into the chow at 0.2%, as previously described^41,42^, and we proceeded with a prospective, controlled, randomized trial in the KL mice. Exposure to ruxolitinib led to an early rise in body weight, which delayed the onset of weight loss and led to a large increase in overall survival (Fig. 4D and E). Ruxolitinib had no effect on tumor burden, as assessed by lung mass (Fig. 4F), but despite not improving skeletal muscle mass, ruxolitinib reversed several features observed in the cachexic muscle (Supp. Fig. 3C and D). For example, JAK/STAT3 activity was suppressed at the protein and transcriptome levels (Supp. Fig. 3E, F and G). A GSEA comparing muscles taken from cachexic mice treated with or without ruxolitinib identified broad reversal of the transcriptomic signatures associated with CACS (Supp. Fig. 3G). Specifically, ruxolitinib increased pathways related to oxidative phosphorylation, ribosomal and proteosome activity, and reduced inflammatory pathways, such as JAK/STAT3, NFKB, chemokines and toll-like receptor signaling. Despite these changes, ruxolitinib had no protective effect on muscle mass (Supp. Fig. 3G).

We noticed that mice with the most weight gain tended to live the longest and performed a correlation analysis of body weight change and survival. The effect of ruxolitinib on overall survival significantly correlated with the amount of weight gained in the initial few weeks of starting the drug (Fig. 5A). Longitudinal body composition analysis showed that mice rapidly gained fat mass after starting ruxolitinib, as opposed to control mice where fat mass was lost over time (Fig 5B). Similar gains in fat mass were observed when ruxolitinib was administered to the KRAS/TP53 (KP) mouse model of lung cancer (Fig. 5C). To study the effects of ruxolitinib before the onset of severe weight loss, we euthanized KL mice eight weeks post induction, and those that had less than 15% of weight loss were categorized as pre-CACS. In agreement with the higher fat mass, plasma leptin levels increased significantly in pre-CACS KL mice (Fig. 5D).

**Figure 5.**
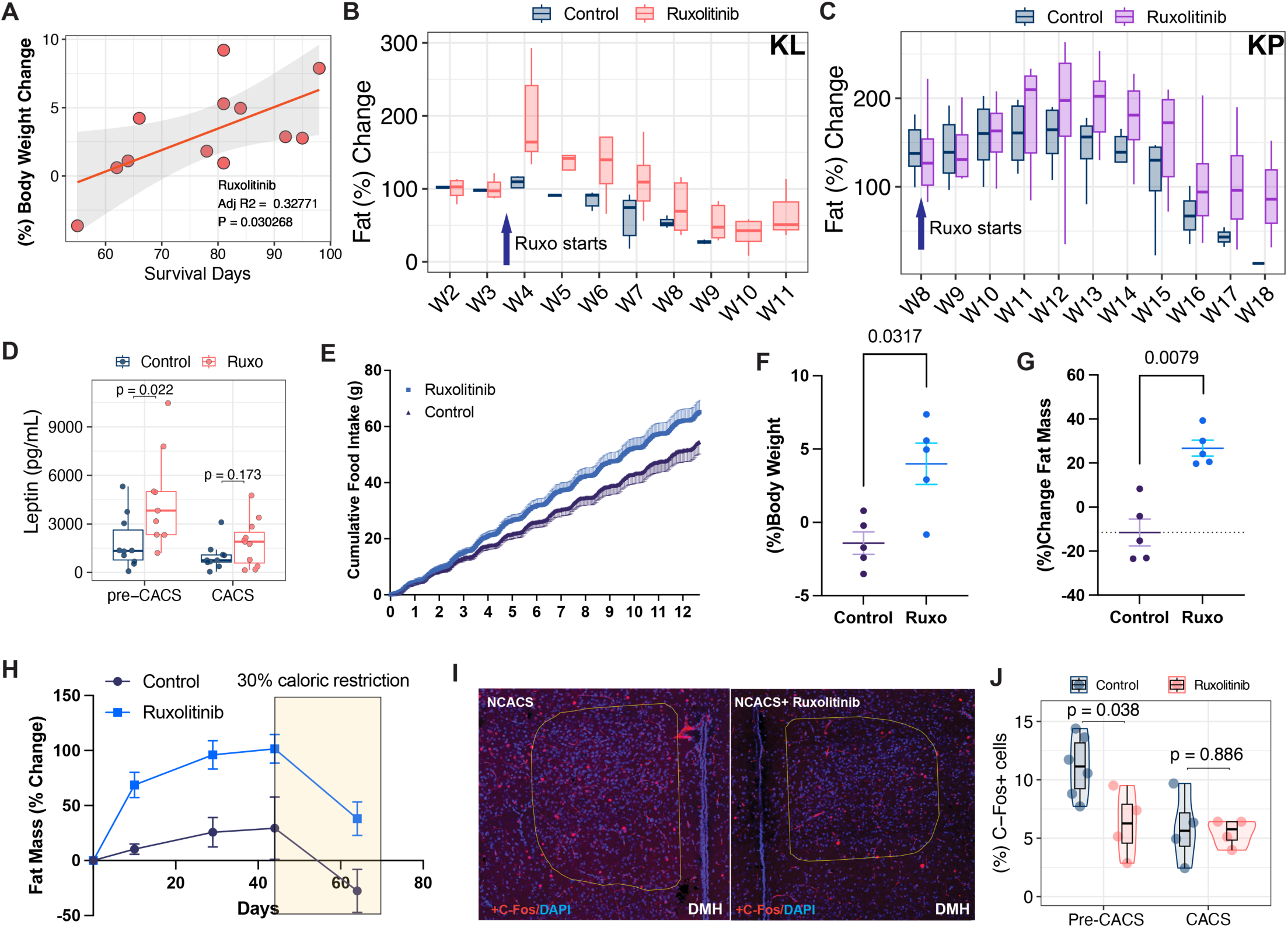
Ruxolitinib prolongs survival by preserving adipose tissue mass. **A.** Correlation analysis of total body weight change between weeks three and six post-tumor induction and survival in days. **B.** Fat mass percentage change in KL mice treated or not with ruxolitinib. **C.** Fat mass percentage change in KP mice treated or not with ruxolitinib. D. Leptin levels in the plasma of KL mice that have not reached 12.5% weight loss (pre-CACS) and CACS, treated or not with ruxolitinib. **E.** Cumulative food intake of non-tumor bearing mice treated or not with ruxolitinib. **F-G.** Percentage change of body weight(F) and fat mass (G) of non-tumor bearing mice treated or not with ruxolitinib for 13 days. **H.** Percentage change in fat mass of non-tumor bearing mice treated or not with ruxolitinib for 45 days and then caloric restricted for 25 days under ruxolitinib treatment or not. **I.** Immunofluorescence staining for p-STAT3 (Tyr 705) on the dorsomedial hypothalamic nucleus (DMH) of KL mice treated or not with ruxolitinib. **J.** Quantification of C-Fos (+) neurons from brains in I. Comparisons in D, F, G and J were performed using Wilcoxon’s test. Individual data points are independent biological replicates unless otherwise stated.

The gain in fat mass from ruxolitinib suggests a net positive energy balance, due to either increased caloric intake or reduced energy expenditure. To assess these options, we treated non-tumor bearing mice with ruxolitinib and found a consistent increase in food intake, body weight and fat mass, with negligible changes in activity or energy expenditure (Fig. 5E, F and G, Supp. Fig. 4A and B). Fat mass may also increase if lipids are retained in the adipose tissue due to defective lipolysis. Because it has been reported that inhibition of JAK/STAT3 directly impairs adipose tissue lipolysis^43^, we treated non-tumor bearing mice with ruxolitinib until they reached weight stability, and then calorie restricted them to promote lipolysis. We observed that both ruxolitinib-treated and control mice lost fat mass at a similar rate (Fig. 5H). These data suggest that ruxolitinib leads to increased body weight and fat by increasing caloric intake, without affecting fat mass mobilization.

Larger amounts of fat mass increase circulating levels of leptin, a crucial regulator of energy homeostasis, caloric intake, and body weight regulation. Inhibition of leptin signaling is known to decrease satiety and increase food intake^44,45^. Since leptin signaling utilizes JAK1/2 and ruxolitinib penetrates the CNS^46^, we hypothesized that the increase in food intake may result from impaired leptin sensing in the hypothalamus. We tested this hypothesis by performing immunofluorescence of c-Fos, a neuronal activity marker, in several hypothalamic areas related to energy homeostasis and food intake. We found c-Fos staining was reduced in the dorsomedial hypothalamic nucleus (DMH) of pre-CACS mice. The DMH is known for its role in energy balance and regulation of circadian feeding mediated by the leptin receptor^47,48^. Similar trends were observed in the arcuate nucleus, ventromedial hypothalamus (VMH) and lateral hypothalamus (LHA), although they did not reach statistical significance (data not shown). These results are in agreement with previous reports linking ruxolitinib-induced weight gain in patients to leptin receptor inhibition^49^.

Lastly, we explored ruxolitinib’s effects in the liver of mice with and without CACS. Ruxolitinib suppressed JAK/STAT3 signaling, reversing several of the transcriptomic signatures previously identified during CACS (Fig.6A, B and C). For example, cachexic mice treated with ruxolitinib showed negative scores for inflammation, complement, JAK/STAT3 signaling and interferon alpha/gamma responses when compared to non-treated (Fig.6C). In agreement, ruxolitinib treatment led to lower levels of APR proteins in the plasma (Fig. 6D). Additionally, ruxolitinib also led to a positive enrichment of pathways related to peroxisome and fatty acid metabolism, including several key PPAR-α target genes required for fatty acid oxidation, and ketogenesis (Fig. 6C, and E). These changes correlated with higher serum levels of beta-hydroxybutyrate in plasma, a marker of hepatic fatty acid oxidation (Fig. 6F). The levels of non-esterified fatty acids and glycerol in the circulation were also higher in ruxolitinib-treated mice (Fig. 6G and H), suggesting that the added fat mass enabled these mice to better maintain the serum supply of lipolytic metabolites.

**Figure 6.**
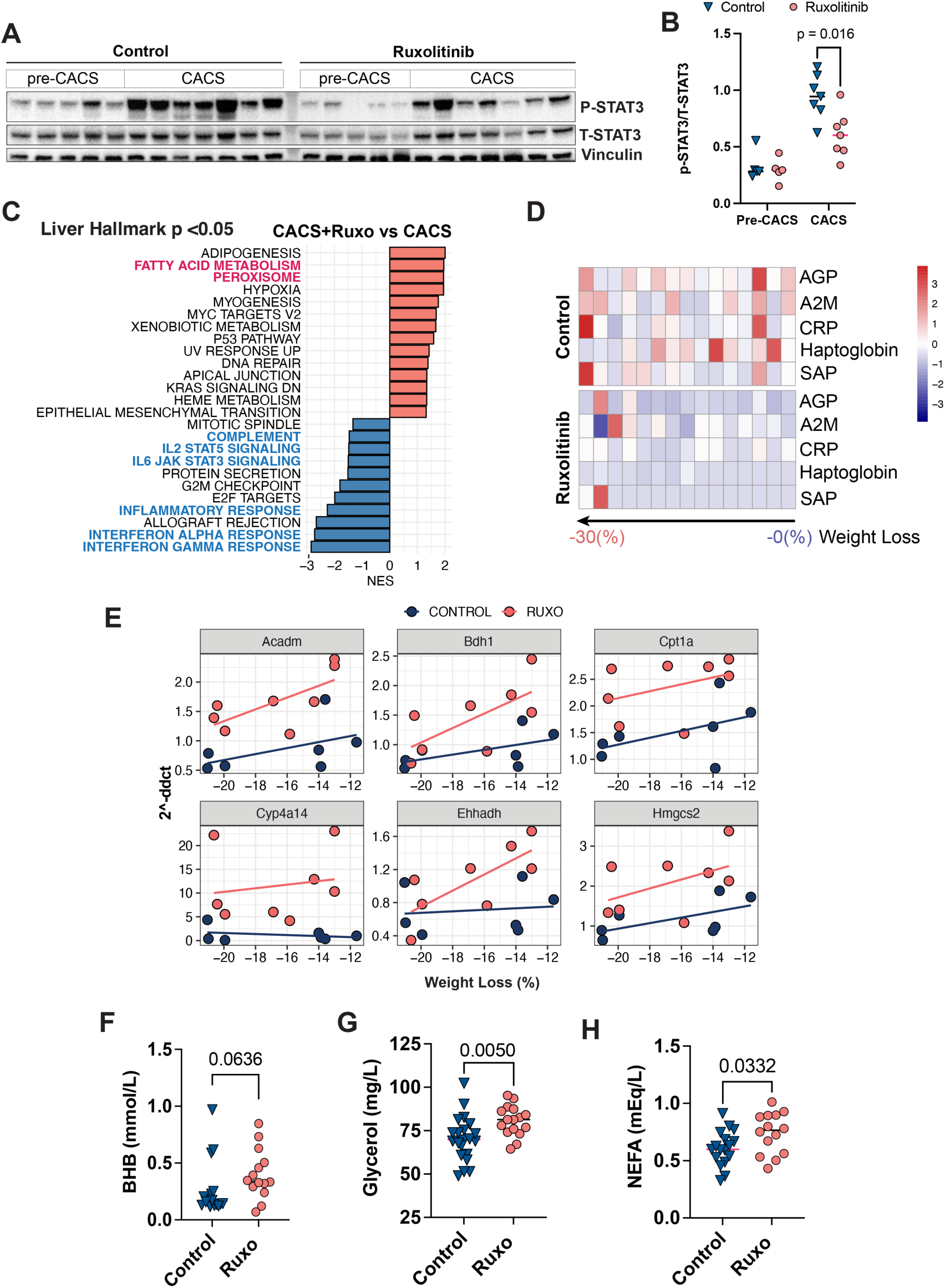
Ruxolitinib reverses PPAR-α inhibition and inflammation in the liver of cachexic mice. **A.** Western blot for p-STAT3 (Tyr705) in the livers of mice treated or not with ruxolitinib **B.** Quantification by densitometry of the bands in A. C. Gene set enrichment analysis (GSEA) of liver transcriptomics from CACS mice comparing the effects of ruxolitinib in the liver of CACS mice treated or not with ruxolitinib. D. Heatmap showing relative plasma levels of acute-phase reactants AGP, A2M, CRP, Haptoglobin and SAP in the plasma of KL mice treated or not with ruxolitinib. E. Expression level of key PPAR-α target genes in the livers of KL mice treated or not with ruxolitinib. **F-H.** Plasma levels before euthanasia of beta-hydroxy butyrate (F), non-esterified fatty acids (G) and glycerol(H) in the plasma of KL mice treated or not with ruxolitinib.

## Discussion

Cachexia is a systemic syndrome with pleotropic effects on various organs, making it difficult to identify causative factors. Across numerous clinical and pre-clinical studies, inflammatory cytokines correlate with cachexia. Here, we find that lung cancer-induced cachexia is associated with a more pro-inflammatory tumor microenvironment, characterized by high expression and secretion of IL-6 family members. Integration of CPTAC and TRACERx datasets confirms that the inflammatory and metabolic changes we observed in KL mice are also evident in patients with lung cancer, supporting the translational relevance of our findings. IL-6 was initially identified as a cachexic factor approximately 30 years ago and continues to be a prominent focus of cachexia research^18,50^. Several studies have reported that IL-6 superfamily members induce atrophy in skeletal muscle and adipose tissue during cachexia^18,51–56^. However, most pre-clinical models of cachexia produce levels of circulating IL-6 that are significantly higher than what is commonly observed in patients with lung cancer^57^. This excessive amount of IL-6 in several preclinical models may lead to overestimating its value as a therapeutic target. Furthermore, the effects of the IL-6 superfamily on cachexia are confounded by their role in tumor progression. The IL-6 superfamily family can contribute to tumor growth^58^, so interventions that block IL-6 may slow cachexia because tumors now grow slower. Recent pre-clinical studies that control food intake and tumor burden suggest that the effects of IL-6 on body weight and tissue mass may be indirect^59,60^. Therefore, the question of whether the IL-6 superfamily contributes to cachexia remains pertinent after all these years.

In KL mice, the abundance of circulating IL-6, LIF, and IL-11 is modest and more representative of that in human patients (Supp. Fig. 1, B,C,D,H and I). In this setting, we were unable to suppress muscle loss by blocking JAK signaling with ruxolitinib. These data are in line with the results from clinical intervention studies targeting IL-6 and challenge the notion that tumor-derived IL-6 superfamily members are sufficient to induce skeletal muscle atrophy in mice and humans with lung cancer^20,21,61^.

What is the role of IL-6 in cachexia?. Our study highlights a set of linked metabolic changes that occur in the liver following activation of JAK signaling. In mice with CACS, the activation of the APR occurs at the expense of PPAR-α-driven fat oxidation pathways^14,36,62^. This phenomenon mirrors the response to bacterial sepsis, where fat oxidation and ketogenesis are also suppressed^63,64^. Suppressing fat oxidation in this setting may shunt adipocyte-derived fatty acids toward reesterification and incorporation into lipoproteins, which are secreted and can neutralize the toxic effects of LPS via direct binding^65^. Furthermore, the loss of oxidative processes in hepatocytes may help create a more reduced environment that facilitates synthesis of triglycerides and APR proteins. It is widely believed that the skeletal muscle supplies the liver with the amino acids necessary to support the APR via JAK signaling^55,66^. Our data from mice treated with ruxolitinib challenge this theory. Ruxolitinib suppressed APR production and maintained fatty acid oxidation in the liver; however, it did not protect muscle mass. We could speculate that despite the reduced production of APR, amino acid demand is still high to supply energetic demands.

Lastly, we find that ruxolitinib delays the onset of cachexia and improves overall survival by increasing food intake, expanding adipose tissue mass, and reducing inflammatory signaling in metabolic organs. While our experimental design does not allow us to determine the relative contribution of each effect, we hypothesize that increased food intake is the primary driver. Prior studies have shown that ruxolitinib inhibits hypothalamic leptin signaling, thereby which increasing appetite and promoting weight gain in both mice and humans^49,67^. In our study, mice that gained the most weight in the first two weeks of treatment, also had the greatest survival benefit. Because adipose tissue serves as a depot for high-density nutrients, these findings suggest that increased fat mass may enhance survival during states of nutrient restriction, such as tumor-induced anorexia. This conclusion aligns with observations in patients with lung cancer, where higher adiposity is associated with longer survival^68^. Notably, previous studies conducted in xenografted colorectal and lung cancer cell lines have suggested that JAK/STAT3 blockade preserves fat by inhibiting lipolysis^12,69^. However, our results show the orexigenic effects of ruxolitinib are most effective in the pre-cachexic state, suggesting that tumor-derived anorectic factors can override canonical hypothalamic circuits. This finding is consistent with the observation that cachexic animals fail to eat despite critically low leptin levels^16^. Whether ruxolitinib can enhance fat mass in patients with lung cancer is currently under investigation (NCT04906746). Our findings suggest that ruxolitinib should be initiated as early as possible in the disease course, before the onset of refractory anorexia and weight loss.

## Methods

### Mouse Model and Tissue collection

For this study, we used the Kras^LSL-G12D/+^; Lkb1^f/f^ mouse model of lung cancer^13^. The mice were kept under a 12-hour light/dark cycle at a temperature of 22 °C, with unrestricted access to rodent chow (PicoLab Rodent 20, 5053, Lab Diet) and water. Lung tumors were induced in adult male and female mice, aged 12-20 weeks, using 2.5×10^7^ plaque-forming units of Adenovirus CMV-Cre (Ad5CMV-Cre) acquired from the University of Iowa Gene Transfer Vector Core (Iowa City, IA). Mice were humanely euthanized using CO2, upon exhibiting either a 30% weight reduction or a deteriorated body condition score of 2 or less. Tissues were dissected, weighed, and flash-frozen in liquid nitrogen and subsequently stored at −80 °C for further use.

### Body Composition and Metabolic Cage Analyses

Fat mass and lean mass were measured using an EchoMRI-100H 2n1 with a horizontal probe configuration (EchoMRI, Houston, TX). For the metabolic cage experiments, mice were single-housed a week prior to the initiation of the experiments. Mice were then placed in the Promethion Metabolic Cage system (Sable Systems, USA), with a 12h light-dark cycles at 22°C. Data from the first 24 h was not used for analysis as it represents an acclimatation period. In the metabolic cages, food and water intake, spontaneous activity, total volume of oxygen consumed, and carbon dioxide liberated are measured every 5 minutes. Mice had access to food and water ad libitum.

### Therapeutic trials

Pacritinib was compounded into the chow at a concentration of 0.3% by TestDiet (St. Louis, MO). Similarly, ruxolitinib was obtained from MedChemExpress (Monmouth Junction, NJ) and compounded to the chow at a concentration of 0.2% as previously published^42^. The placebo control groups either received IP injections of vehicle solution or normal chow. There was no blinding and therapeutic trials were carried on male mice. The primary outcome was overall survival unless otherwise stated. Secondary outcomes included body weight changes, body composition measures (fat/lean mass), tumor burden (lung mass), food intake, and organ weights. For pacritinib and ruxolitinib, NCACS mice were excluded from survival or weight loss analysis to allow for evaluating the treatment effect on cachexic mice only.

### Cytokine Analysis

Immediately following euthanasia, mouse blood was obtained via cardiac puncture, centrifuged at 10,000×g for 10 minutes at 4 °C, and the serum was then frozen at −20 °C for subsequent testing. During necropsy, some lungs were excised for tumor isolation immediately frozen in liquid nitrogen, and stored at -80°C. Tumors were lysed to measure cytokines using a specialized lysis buffer (200mM EDTA, 5M NaCl, 1% NP-40, 0.5% Triton X-100, 10% Glycerol, and 1M Tris-base), and then diluted to a concentration of 1 mg/ml with PBS. Eve Technologies (Calgary, AB) conducted the initial cytokine screening for both serum and tumor lysates. Mouse cytokines not included in Eve technologies panel were evaluated using Mouse Luminex Discovery Assay from R&D (IL-33, IL-6, LIF, Tweak, IL-27, IL6Ra and FGF21) or Milliplex Mouse Cytokine/Chemokine Magnetic Bead Panel II (IL-11). AGP, A2M, CRP, Haptoglobin and SAP, in mouse serum were measured with Milliplex MAP Mouse Acute Phase Magnetic Bead Panel 2. Cytokines in human plasma were measured using Human Magnetic Luminex Assays from R&D.

### Western Blots and Antibodies

Gastrocnemius, TA, and liver were lysed using a tissue lysis buffer containing 50 mM Tris·HCl (pH 7.4), 150 mM NaCl, 1 mM EDTA, 10% glycerol, 1% Nonidet P-40, 0.5% Triton X-100, and protease and phosphatase inhibitor (Pierce, A32959). Samples were then denaturalized at 70°C for 20 min and loaded onto 4–12% NuPAGE Bis-Tris gels at a concentration of 30-50 μg and transferred to 0.45-μm PVDF membranes with wet transfer cells (Bio-Rad Laboratories). After 1 h of blocking with Tris-buffered saline with 0.1%, Tween 20 containing 5% BSA (TBST), membranes were incubated overnight at 4 °C with antibodies against p-STAT3 (Tyr705, CST #9145), Total STAT3 (Sc-8019), and vinculin (CST #4650). Membranes were incubated with either anti-mouse (CST# 7076) or anti-rabbit (CST# 7074) secondary IgG-HRP linked antibodies. Membranes were revealed using either HyBlot CL Autoradiography Film (Denville Scientific) with SuperSignal Western Blot enhancer solution (Thermo Fisher), or with a ChemiDoc (BioRad).

### ELISA and other colorimetric assays

LPS concentration in mouse tumor lysates was measured by ELISA (LSBio, LS-F17912-1), per the manufacturer’s instructions. LPS concentration in mouse liver lysates was done with ENDOLISA (Biomerieux, Cat. No. 609033), per the manufacturer’s instructions. Prostaglandin E2 was measured in mouse tumor lysates with an ELISA from R&D (cat. No. KGE004B), per the manufacturer’s instructions. Alanine Aminotransferase Activity Assay Kit (Sigma, MAK052). Glycerol Colorimetric Assay Kit was purchased from Cayman (10010755). B-Hydroxybutyrate Liquicolor Assay, Stanbio 2440. NEFA was purchased from Fujifilm (997-76491, 276-76491,993-35191, 991-34891, 995-34791, 999-34691).

### Histological and Immunohistochemical Procedures

Mice tissues were fixed in 4% paraformaldehyde (PFA) overnight, followed by preservation in 70% ethanol. Paraffin-embedding, sectioning (4μm), H&E staining and slide scanning were done at Histowiz (Long Island City, NYC, USA). Slides for IHC and IF were deparaffinized in Histoclear (National Diagnostics), rehydrated, and subjected to antigen unmasking using either citrate buffer (10 mM Citrate, 0.05% Tween 20, pH 6.0) for F4-80 CST #70076 or Tris-EDTA buffer, pH 9.0 (10 mM Tris Base, 1mM EDTA, 0.05% tween 20) for p-STAT3, CST #9145 in a pressure cooker for 13 minutes. To inhibit endogenous peroxidase, a 3% hydrogen peroxide solution in PBS was used for 5 minutes. Blocking was achieved with 5% rabbit or donkey serum. The slides were then incubated with primary antibodies overnight. Following steps included the application of secondary antibodies such as Goat Anti-Rabbit IgG Antibody (H+L), Biotinylated, R.T.U. (BP-9100-50, Vector) or HRP Goat Anti-Rat IgG Polymer Detection Kit, Peroxidase (MP-7404-50, Vector), avidin/biotin reagents (PK-4100) and DAB peroxidase substrate from Vector Laboratories (SK-4100). Imaging was done using either an Aperio AT2 (Leica Biosystems) or a Zeiss Axioscope Imager at Histowiz. IHC quantifications were automated using whole slide scans and QuPath software (0.4.2)^70^. Brains were fixed with 4% PFA for 18 hs and sectioned into free-floating sections by Neuroscience Associates (Tennessee, USA). C-FOS immunofluorescence was done using sc-166940 antibody in free-floating sections as previously described^71^. Slides were imaged using a Zeiss 880 Confocal microscope, and quantification was done with the Qupath extension ABBA^72^.

### Patient Sample Collection

Plasma samples were collected in K2 EDTA tubes from lung cancer patients at New York Presbyterian Hospital/Weill Cornell Medical College, in compliance with an IRB-approved protocol (IRB#19-11021135). Plasma samples (K2 EDTA) from healthy individuals were sourced from Innovative Research (https://www.innov-research.com/). Demographics of patients and healthy controls are detailed in Table.1.

### Osmotic pumps implantation

Osmotic pumps were purchased from Alzet (model 2001). Implantation of osmotic pumps was performed following the manufacturer’s guidelines. Briefly, after anesthetizing the animal, the skin over the mid-scapular region is shaved and washed. Next, the subcutaneous tissue is dissected with a hemostat to create a pocket large enough to accommodate the pump. The pump filled with either IL-6 (cat. No 575708) or PBS is then inserted into the pocket, with the delivery portal oriented caudally. Lastly, the wound is closed with wound clips. The rate of IL-6 infusion was 1µL/hour at a concentration of 0.3 mg/mL.

### Drugs

Ruxolitinib phosphate (HY-50858), was purchased from MedChemExpress. Pacritinib was kindly provided by CTI Biopharma.

### Real-Time PCR

RNA was extracted from primary mouse hepatocytes and tissues using TRIzol (Thermo Fisher) and in the case of tissues, it was further purified using the RNeasy kit (Qiagen). cDNA synthesis was done with SuperScript VILO Master Mix (Thermo Fisher). PPAR-alpha target genes in the liver were quantified with Applied Biosystems TaqMan Gene Expression Assays using the following assays: Rplp0 (Mm00725448_s1), Actb (Mm00607939_s1), Ehhadh (Mm00619685_m1), Cyp4a14 (Mm00484135_m1), Fgf21 (Mm07297622_g1), Pdk4 (Mm01166879_m1). 16S rRNA qPCR was done following a previously published protocol^73^. mRNA levels were quantified using the 2^-ΔΔCt method.

### Bulk RNA-sequencing

Tumor, liver and gastrocnemius RNA-sequencing of the KL mice that were previously published are available at the GEO Database (GSE107470, GSE165856)^14,74^. New RNAseq datasets exlclusive to this manuscript are also available at the GEO Database (GSE308735). Liver and gastrocnemius RNA-sequencing of the Kras^G12D/+;^ Lkb1^f/f^, treated or not with ruxolitinib were done by GENEWIZ, LLC. Azenta US, Inc. (South Plainfield, NJ, USA). Briefly, RNA sequencing library was prepared using the NEBNext Ultra II RNA Library Prep Kit for Illumina using manufacturer’s instructions (New England Biolabs, Ipswich, MA, USA). The sequencing library was validated on the Agilent TapeStation (Agilent Technologies, Palo Alto, CA, USA), and quantified by using Qubit 2.0 Fluorometer (ThermoFisher Scientific, Waltham, MA, USA) as well as by quantitative PCR (KAPA Biosystems, Wilmington, MA, USA). Reads were aligned using Galaxy (https://usegalaxy.org/). Briefly, paired-end fastq files were processed with Cutadapt for adapter removal. Raw reads were mapped using STAR (2-pass mapping) with the GENCODE M25 annotation on the mm10 genome assembly. Counts were obtained with featureCounts, and quality control was performed with FastQC and MultiQC. Differential expression analysis was done in R Studio (2023.03.0+386), with R (4.2.3) and DESeq2 (v.1.38.3). Gene set enrichment analysis (GSEA) was performed using the GSEA (v4.3.2) JAVA-based application and mouse Molecular Signatures Databases published by the BROAD Institute and the UC San Diego (https://www.gsea-msigdb.org/gsea/index.jsp). Mouse KEGG pathways were obtained from easyGSEA (https://tau.cmmt.ubc.ca/). Tumor pathways analysis in Fig. 1E was done with Qiagen’s Ingenuity Pathway Analysis (IPA, v. 01-21-03). Heatmaps were done using heatmap (v. 1.0.12) and volcano plots with EnhancedVolcano (v. 1.16.0). GSVA was performed using the GSVA R package (v.1.46.0). Venn diagrams were done using the R package given (v. 0.1.10). TracerX and CPTAC datasets are publicly available.

### Software and Statistical Analysis

Data analysis was primarily conducted using RStudio (2023.03.0+386) on R (v. 4.2.3) using the statistical packages ggpubr (0.6.0), rstatix (v.0.7.2) and ggplot2 (v. 3.4.4) and GraphPad Prism 9. Mouse Kaplan-Meier survival curves were compared using Log-Rank tests within the R packages “survival” and “surviminer” which also utilizes “maxstat” for dichotomizing continuous data using maximally selected rank statistics. Kaplan-Meier survival curves for the Pan-Cancer combined caBIG, GEO and TCGA databases were plotted and analyzed using the default parameters in (https://kmplot.com/analysis/).

## Data Availability Statement

All data generated in this study is publicly available. The newly generated bulk RNA-Seq has been deposited in the GEO Database (GSE308735). Previously published RNA-Seq data are accessible at GSE107470 and GSE165856. No original code was generated specifically for this paper.

---

**Table 1.**
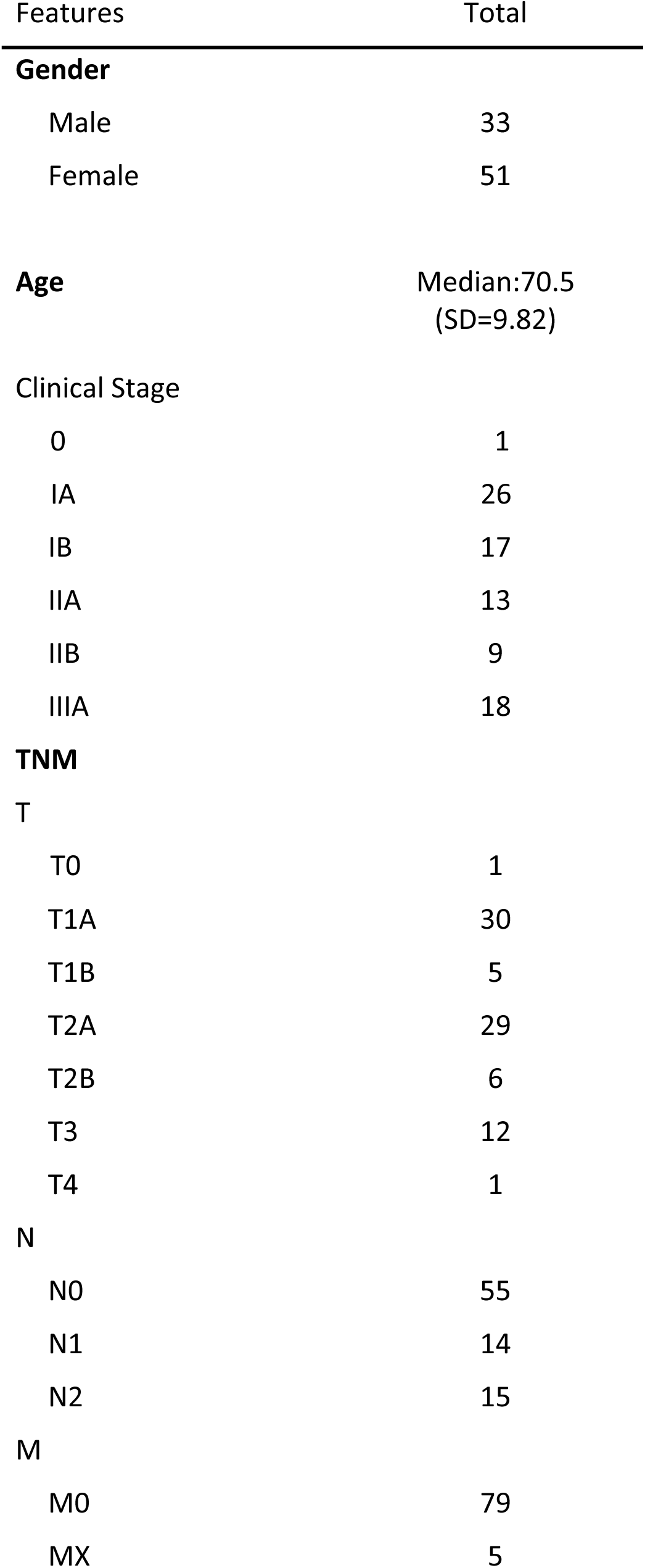
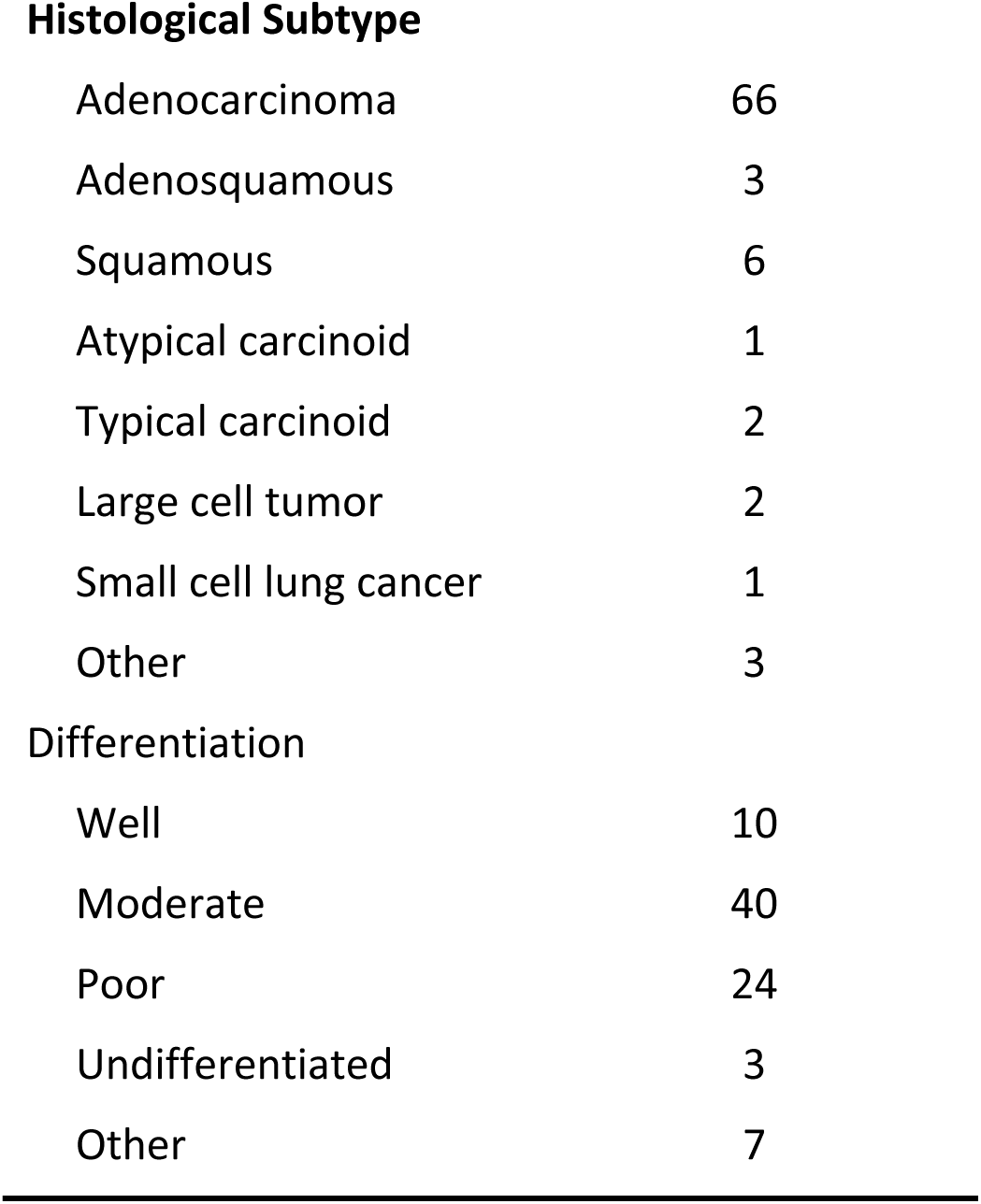
Demographic and clinical details of patients with lung cancer in this study.

**Table 2.** Demographics of healthy control plasma donors used for this study.

| Features | Total |
| --- | --- |
| Gender |  |
| Male | 17 |
| Female | 18 |
| Age | Median:37(SD=16.42) |

## Conflict of interest statement

M.D.G. holds equity in Faeth Therapeutics and Skye Biosciences; reports consulting or advisory roles with Almac Discovery, Faeth Therapeutics, Genentech Inc., Scorpion Therapeutics, Skye Biosciences, and Third Arc Bio, Inc.; honoraria from Novartis AG, Pfizer Inc., Genentech Inc., patents, royalties, and other intellectual property with Weill Cornell Medicine and Faeth Therapeutics. E.D. reports intellectual property with Weill Cornell Medicine. All other authors report no conflicts.

## Funding statement

This work was supported by the 2020 AACR-AstraZeneca Lung Cancer Research Fellowship, 20-40-12-DANT (E.D.), a grant from the Starr Cancer Consortium (D.J.P., O.E., B.M.S.), and was delivered as part of the CANCAN team supported by the Cancer Grand Challenges partnership funded by Cancer Research UK (CGCATF-2021/100022) and the National Cancer Institute (1 OT2 CA278685-01).

## Author

E.D., A.M., T.A., M.A., J.K., T.P., S.R., I.N., A.L.Q., E.C.L., M.G.J., S. L.,J.G., B.E. and M. F. performed experiments; E.D, M.D.G, designed the project; B.E., I.G.,T.D.,M.J.H., T.J., E.W., M.M., N.A, provided technical support; E.D., A.M., T.A., M.A., T.P., S.R., M.N., I.N., A.L.Q. T.L. and M.F. analyzed the data. E.D., M.D.G., wrote the paper. All authors read, edited, and approved the manuscript.

## Acknowledgements

We gratefully acknowledge the patients and relatives who participated in the TRACERx study and the TRACERx consortium. Pacritinib was graciously provided by CTI BioPharma.

---

**Supp. Fig. 1.**
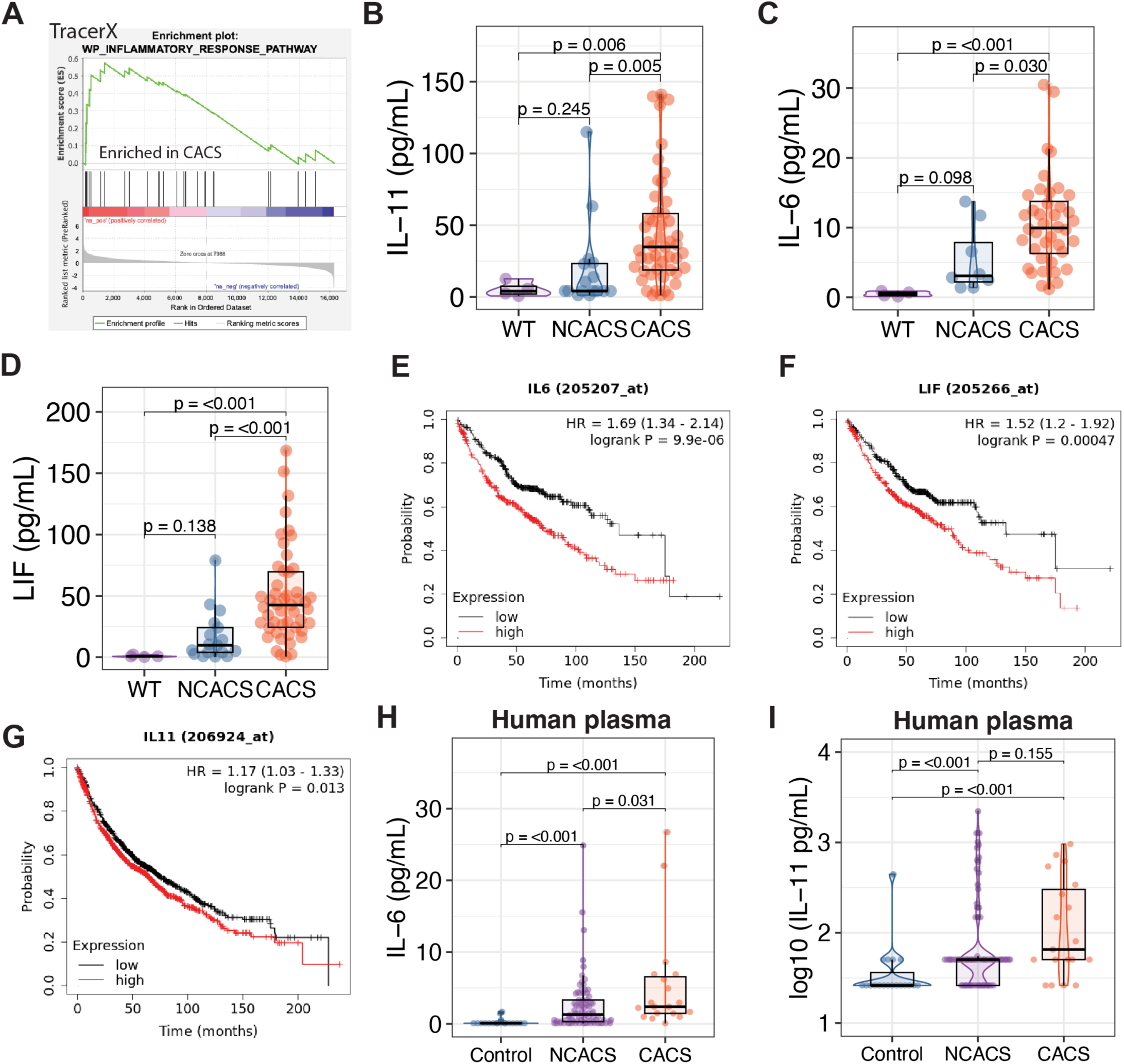
IL-6 family members are associated with an unfavorable prognosis in human lung cancer. **A.** GSEA comparing CACS to NCACS in tumor transcriptomics from the TracerX cohort, showing increased inflammatory response in the tumors of patients with cachexia. B-D. Circulating levels of IL-11 (B), IL-6(C) and LIF(D) in non-tumor bearing mice (WT), NCACS and CACS mice. **B-D.** Kaplan-Meier plots showing survival probability of patients with lung adenocarcinoma stratified by high or low tumor expression of IL-6, LIF and IL-11. **E and F.** Protein concentration measured by Luminex of IL-6(E) and IL-11(F) in the plasma of lung cancer patients with and without cachexia and control plasma samples. Comparisons in E and F were done with the Kruskal-Wallis test, followed by Dunn’s test. Individual data points are independent biological replicates unless otherwise stated.

**Supp. Fig. 2.**
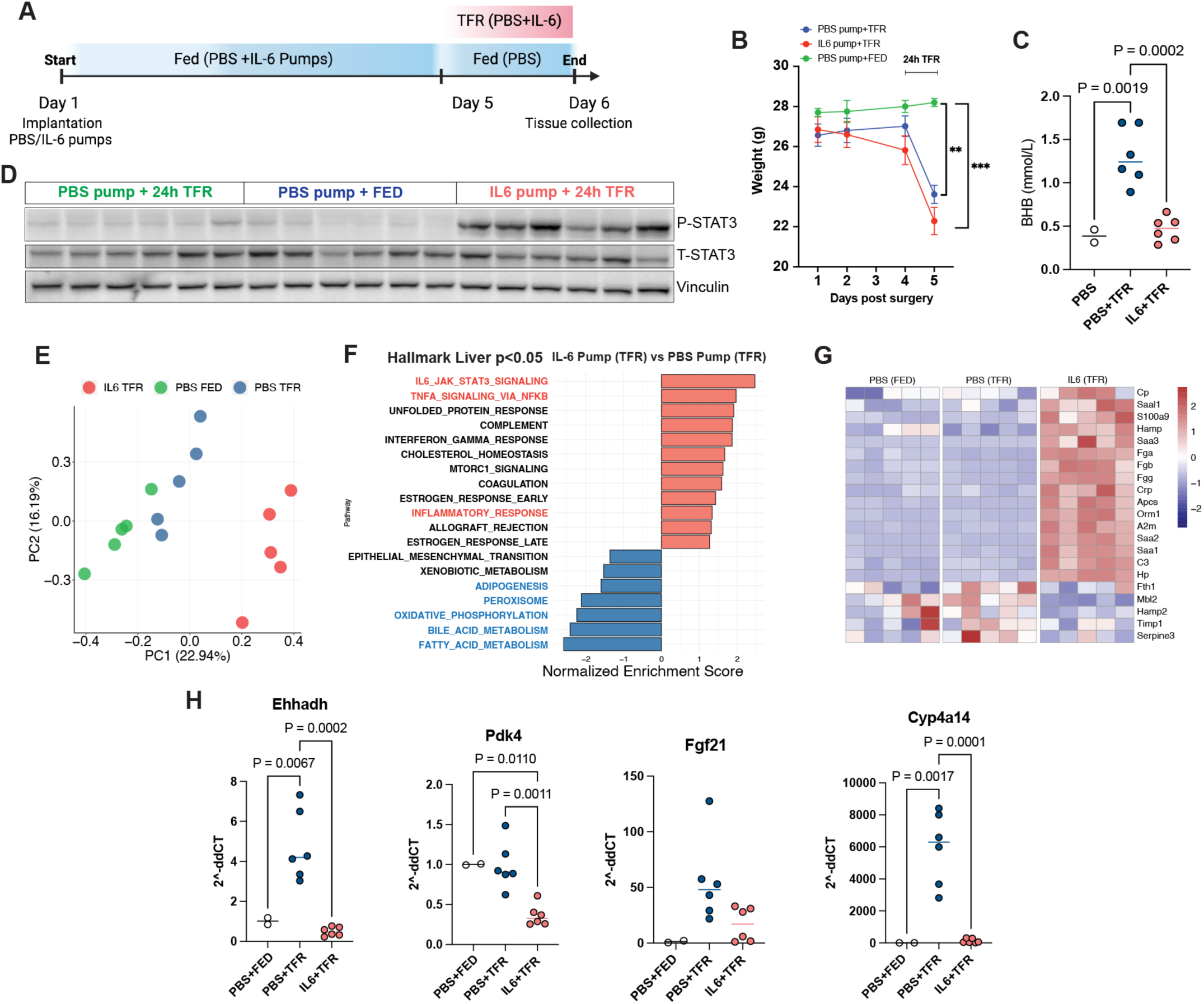
IL-6 reproduces the liver phenotype of the KL CACS mice. **A.** Experimental design for non-tumor bearing mice implanted with IL-6 or PBS secreting pumps (“Fed” is unrestricted access to food, “TFR” is total food restriction for 18 hs). **B.** Whole-body weight of mice implanted with IL-6 secreting pumps and total food restriction for 24hs (TFR, red), mice with PBS pumps and 24hs TFR (blue), and PBS pumps with ad-libitum food (green). **C.** Beta-hydroxybutyrate (BHB) measured in the serum of the mice in A/B. **D.** WB for p-STAT3 (Tyr705), Total STAT3 and Vinculin in the liver of mice implanted with PBS or IL-6 pumps **E.** Unbiased principal component analysis (PCA) of liver transcriptomics from WT (non-tumor bearing) mice implanted with IL-6 secreting pumps and total food restriction for 24hs (TFR, red), mice with PBS pumps and 24hs TFR (blue), and with PBS pumps but with ad-libitum food (green). **F.** GSEA of liver transcriptomics comparing Hallmark pathways between mice implanted with IL-6 or PBS-secreting pumps and TFR for 24hs. **G.** Heatmap of APR-related genes in the liver transcriptomics of mice implanted with PBS or IL-6 pumps. **H.** qPCR for PPAR-α target genes in livers of mice implanted with PBS or IL-6 pumps. Comparisons in C and H were performed with one-way ANOVA followed by Tukey’s multiple comparisons test. Comparisons in J were done using a two-tailed Student’s t-test. Comparisons in B were done with two-way ANOVA followed by Tukey’s multiple comparisons test (**=0.0008, ***=<0.0001). Individual data points are independent biological replicates unless otherwise stated. Model in (A) was made with Biorender.com.

**Supp. Fig. 3.**
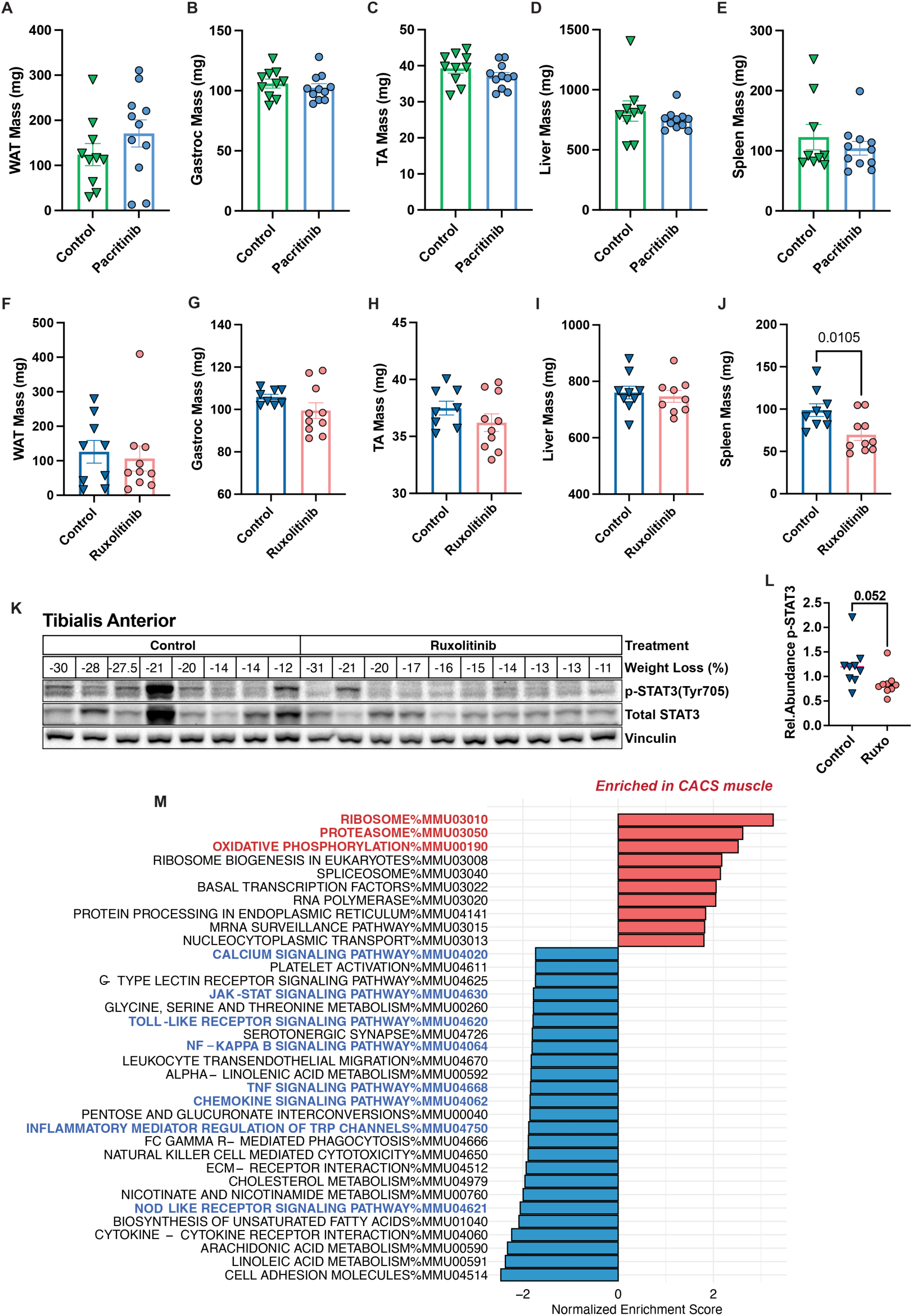
Effects of JAK/STAT3 inhibition in peripheral tissues. **A-B.** Averaged tissue weights from KL mice at endpoint treated or not with pacritinib. **C-D.** Averaged tissue weights from KL mice at endpoint treated or not with ruxolitinib. **E.** WB for p-STAT3 (Tyr705), Total STAT3 and Vinculin in the tibialis anterior of KL male mice treated or not with ruxolitinib. **F**. Quantification by densitometry of the p-STAT3 WB shown in (K). **G.** GSEA using the Kyoto Encyclopedia of Genes and Genomes (KEGG) database to compare muscle from cachexic mice treated or not with ruxolitinib (q-val <0.001). Comparisons between control and pacritinib or ruxolitinib-treated mice in A-J and L were done using a two-tailed Student’s t-test. Individual data points are independent biological replicates unless otherwise stated.

**Supp. Fig. 4.**
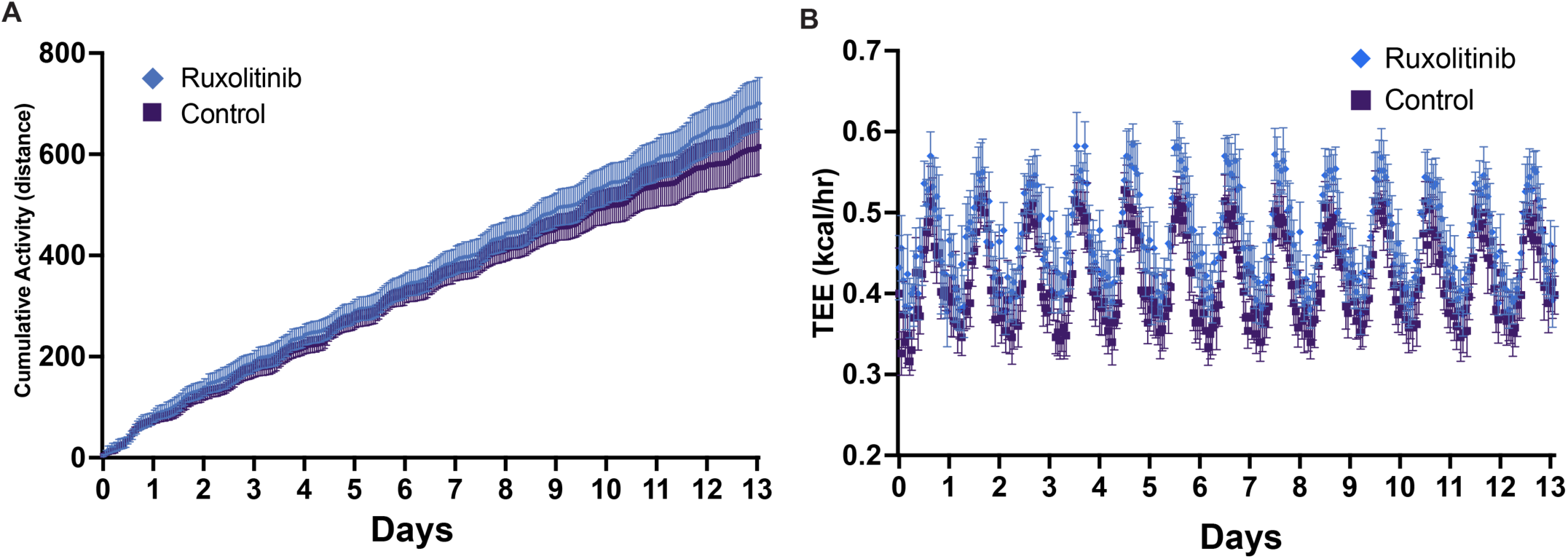
Ruxolitinib does not affect activity or energy expenditure in non-tumor bearing mice. **A.** Cumulative activity of non-tumor bearing mice treated or not with ruxolitinib. B. Total energy expenditure of non-tumor bearing mice treated or not with ruxolitinib.

